# Enhanced data collection in the Canadian Arctic for seabird bycatch information yields highly variable results

**DOI:** 10.1101/2024.11.29.626127

**Authors:** Jennifer F. Provencher, André Morrill, Mark L. Mallory

## Abstract

Incidental catch of seabirds (bycatch) in fisheries has been identified as a major threat to the conservation of seabird populations globally. Acquiring accurate, detailed data on seabird bycatch is an ongoing challenge to effective integrated ecosystem management of commercial fisheries. This is especially true in the Arctic region where different countries have highly variable reporting and data systems to track and understand seabird bycatch in fisheries. To collect detailed data on seabird bycatch in the Greenland halibut (*Reinhardtius hippoglossoides*) fishery in northern Canada, we applied two methods that asked for more information than standardly reported in the fishery as a voluntary effort with industry partners. We found that the amount of bird bycatch reported in both enhanced datasheets completed by at-sea observers (ASO) and carcass collections yielded different results when compared to the typical seabird bycatch reporting in the fisheries ASO database. Across three years of data collection (2016, 2018, and 2019), the number of seabirds reported using the enhanced data collection methods were 0.5-11-fold the number from typical ASO database values. In a fourth year of observations (2023), enhanced datasheets from the ASO reported no bycaught fulmars (gulls and terns were reported), but the accompanying photographs showed northern fulmars as bycatch, further supporting that seabird identification is hampering accurate reporting. We then used these data to model how the differences between data sources may fluctuate across years. These large discrepancies between the methods highlight the challenges with obtaining accurate and precise seabird bycatch data needed to implement a meaningful ecosystem approach to the management of the fisheries. Future modelling efforts need to take these differing data sources and variability into account to fully understand the potential population level impacts of fisheries on seabird populations.

## Introduction

Seabirds accidentally get entangled, or caught, in fishing gear as part of fishery activities (Pott and Wiedenfeld 2017). Incidental catch of seabirds (i.e., when the birds are not targeted, but nonetheless are caught in the gear as part of the regular fishing operations) is termed seabird bycatch. Typically, seabirds caught in fishing gear die (Pott and Wiedenfeld 2017). Seabird bycatch has been recognized as one of the largest threats to seabirds globally, with hundreds of thousands of birds being lethally caught in fisheries each year (Anderson et al. 2011; Žydelis et al. 2013; Dias et al. 2019). Much research and international concern has been focused on bycatch in mid- or lower latitudes, as well as albatrosses and petrels in the Pacific and Southern oceans (e.g., Tuck et al. 2003; Petersen et al. 2009; Phillips 2013; Jiménez et al. 2014), but increasing attention is now focused on the North Atlantic and even more northern waters (e.g., Merkel 2011; Regular et al. 2013; Christensen-Dalsgaard et al. 2019; Mallory et al. 2022). While fisheries in Arctic Canada are relatively small with only a few license holders compared to other regions, the seabird bycatch rates on a per boat basis may be much higher than in Atlantic Canada (Hedd et al. 2016). Importantly, during recent work examining seabird bycatch in Arctic Canada within a cumulative effects context, fisheries have been identified as one of the largest negative impacts on the local seabird population (Rooney et al. 2023).

Although multiple Arctic marine bird species are susceptible to fisheries bycatch, there is a critical requirement for detailed information on seabird bycatch in fisheries from Arctic Canada because of the potential negative effects from fisheries on northern fulmar (*Fulmarus glacialis*) numbers (Anderson et al. 2018). Previous reports have highlighted that northern fulmars are susceptible to being incidentally caught by fisheries in the Arctic region during the open water period (Hedd et al. 2016; Anderson et al. 2018). Recent colony assessments of northern fulmars in Arctic Canada have also shown that the local numbers are declining, and that colony size may be 75% lower than previous estimates (Anderson et al. 2018; Mallory et al. 2020). Population modelling of northern fulmars in relation to fisheries bycatch in Arctic Canada has also shown that while age structure and fishing level can potentially influence the trajectory of northern fulmar population over time, the size of the Arctic population of fulmars from which fisheries were removing individuals has the greatest impact on whether the bycatch was having significant negative impacts (Anderson et al. 2018). Thus, the only way to fully understand the current potential impact regional fisheries may have on Arctic-breeding northern fulmar numbers in Canada is to assess both the fisheries bycatch levels and fulmar population counts in demographic modelling exercises.

In Arctic Canada, there are currently only a few fisheries that are commercially active in the Baffin Bay and Davis Strait region (North Atlantic Fishery Organization (NAFO) Subarea 0 (SA0) Divisions 0A and 0B (0AB)). The Greenland halibut (*Reinhardtius hippoglossoides*) fishery takes place in most years from 1 May to 31 December. While the fishery in this region changes from year to year, in the most recent report on the fishery (2023), industry reported that there were active gillnet and active trawl licences operating in the fishery, with a total of five vessels operating under the gillnet licences and four vessels operating under the trawl licences (Knapman et al. 2023).

Previous work has reported on seabird bycatch in the NAFO 0AB Greenland halibut fishery using the at-sea observer (ASO) data accessed via the Department of Fisheries and Oceans Canada (DFO) (Hedd et al. 2016; Anderson et al. 2018). In this fishery, the ASO companies are third-party groups hired by the industry and are responsible for collecting data on both the target species and any incidentally bycaught species to provide both industry (the quota holders) and the regulator (DFO) with the data to manage the fishery. The requirement and level of observer coverage in the fishery is a condition of the fishery license set by DFO and varies from 20% (in 0B) to 100% (in 0A) coverage targets based on gear and season and management area (DFO 2007; Knapman et al. 2023). As DFO has a responsibility for reporting on incidental catch of all species (including seabirds) in commercial fisheries managed by DFO (DFO 2007, 2019a,b), seabirds are reported by fisheries ASO alongside the target fishery and any incidentally caught fish and marine mammals (Hedd et al. 2016; Anderson et al. 2018).

As part of our collaborative work with the Nunavut Fisheries Association (NFA), previously known as the Nunavut Offshore Allocation Holders Association (NOAHA), and DFO we developed this project to better inform estimates of incidental catch of seabirds in the fishery. This included two ways to collect additional information about seabird bycatch via enhanced datasheets and carcass collections. The aim was to provide detailed seabird bycatch data specific to the 0AB Greenland Halibut fishery to help assess the potential impacts the fishery may have on seabird populations in the region.

In this paper, we outline methods and present data on three different methods to collect information and samples on seabird bycatch in the Greenland halibut fishery in NAFO 0AB. Due to the known challenges with seabird bycatch identification in the field, we expected that while birds of unknown species were submitted, the vast majority, if not all the birds submitted, would be fulmars. In order to compare the two enhanced methods with traditional ASO data collection, we compared the total fulmar numbers reported here to those reported elsewhere directly from the ASO database accessed from DFO (Knapman et al. 2023). While we predicted that reporting via these enhanced seabird bycatch methods would yield more details about the potential interactions with gear and improve species identification, we expected that the overall numbers of bycaught birds reported for a season would be lower than those in the ASO database, as we assumed that not all observers would participate in the voluntary enhanced data collection methods employed in this study. Importantly, this additional enhanced data collection was completed on a voluntary basis by ASO who were already on the vessels, and who were primarily responsible for reporting in the traditional bycatch data collection methods.

## Methods

### Recording of seabird bycatch by at-sea observers (ASO)

Briefly, data on seabird bycatch in fisheries in Canada comes from ASOs that are stationed on active fishery vessels. They collect information on both the target fisheries and any bycatch species that are brought on board. As outlined in previous reports, the ASO-compiled database for a fishery may be accessed via DFO (Hedd et al. 2016; Anderson et al. 2018). Importantly, the ASOs in the 0AB fisheries report in kilograms of species of birds. This is a result of the ASOs only reporting on dead birds that are brought on board in the fisheries gear being used. We used summaries of these data from other sources to compare to our enhanced methods findings (Knapman et al. 2023).

### Enhanced seabird bycatch data sheets

In 2016, 2017, 2018 and 2023 we worked with DFO, the third-party ASO companies in eastern Canada, and industry partner NFA to deploy enhanced seabird bycatch datasheets (SI Figure A). These enhanced seabird bycatch datasheets were modelled after the enhanced marine mammal datasheets that DFO regularly employs in fisheries to improve data collection on species of special concern. The enhanced seabird bycatch datasheets collected the number and species of dead birds captured in the gear, which the ASO report in their standard data methods collections, but also asked the ASO to indicate if birds were caught in gear and released alive, or sustained a major injury in the gear but were not released or escaped the gear without being brought on board (birds which are not reported in the standard ASO data in Canada) (Anderson et al. 2018). The gear type and use of bait was also reported on the enhanced datasheets, to provide additional insights on how bird-gear interactions were taking place. These enhanced seabird datasheets were provided by Acadia University in hard copy form as part of the larger sampling kits (Figure 1A; expanded on more in the following section). In the case of the 2023 enhanced seabird bycatch, datasheets were also submitted with associated photographs of the birds reported as indicated on the reporting sheets.

**Figure 1.**
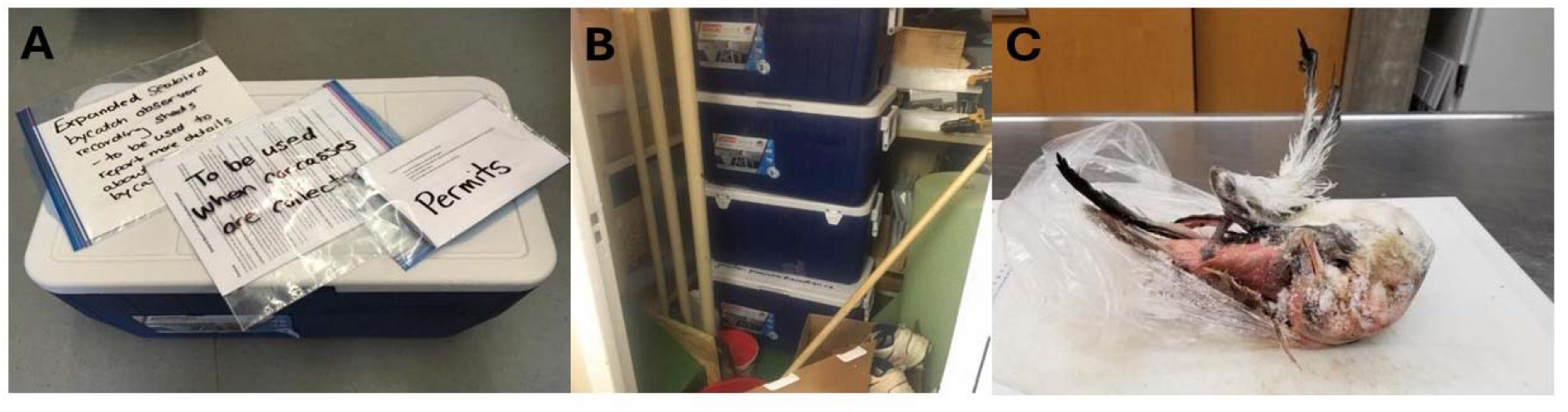
A – sampling kits prepared by Acadia University, including the enhanced seabird bycatch data sheets for at-sea observers to complete, carcass collections, and the necessary permits. B – a picture of the coolers on a fishing vessel shared by the Nunavut Fisheries Association. C – a fulmar carcass that was submitted to ECCC as part of the sampling program in 2018.

### Carcass collections

In 2016, 2017, and 2018 we also worked with DFO and NFA to deploy sampling kits to vessels to collect fulmars for research purposes (e.g., genetic colony assignment and plastic ingestion). Sampling kits were prepared by Acadia University and distributed directly to the vessels (Figure 1A). The collection effort of carcasses was unequal across the years of the study based on partners’ ability and capacity to contribute to the project. The quota holders in Division 0AB agreed to carry the sampling kits and collect bycaught birds, including northern fulmars and other specimens with unknown identification (Figure 1B).

In addition to gathering current information on incidental marine bird bycatch, we had two main purposes for the collections. First, the collection of carcasses was requested by the fishing industry partners so we could conduct a colony assignment genetics study, which was part of the work identified by Environment and Climate Change Canada (ECCC) and fishing partners as necessary to better understand what breeding populations of northern fulmars were being affected by bycatch in 0AB (Colston-Nepali et al. 2020). All carcasses submitted to ECCC were sampled for breast muscle tissue and submitted to partners who carried out the genetics study (Colston-Nepali et al. 2020). A second goal of the project was to also examine the ingested plastic pollution by bycaught fulmars as part of ECCC’s ongoing plastic pollution monitoring program (ECCC 2021).

### Data treatment

We report on bycatch species, collection month, and gear type when that information was available. All data presented were from the Greenland halibut fishery in Division 0AB as this was the fishery of interest for the study. We omit details on ASO IDs, vessel names, trip number, set numbers, and specific dates to ensure that the data are anonymized to the fleet level. We report generally on seabird bycatch from 0AB, and not the separate Divisions of 0A and 0B because some of the reported data did not specify which division within 0AB the data was coming from, and we lacked more detailed metadata to cross check and complete the data blanks in the reports. To compare the quality and quantity of the data collected using the different seabird bycatch data collection tools (enhanced datasheets and carcass collections), we compare the numbers reported here with data previously reported ASO data reported in Knapman et al. (2023).

### Simulation

We used a simulation modeling approach to explore how understood differences in the reporting of fulmar bycatch between the ASO database and the enhanced datasheets could drive variation in observable and documented bycatch across years. These simulations served an illustrative purpose, demonstrating the consequences of our assumptions about the reported bycatch data-generating process, even when assuming low underlying levels of mis- or under-reported bycatch which were still consistent with observations. Our aim was not to make further inferences about reported fulmar bycatch rates beyond what was concluded from our observations, as the simulation output could only corroborate those assumptions. Instead, we sought to visualize the potential contrasts and long-term variation in observable and documented bycatch that arose naturally from our interpretations.

True fulmar bycatch levels in the NAFO 0AB Greenland halibut fishery are never observed and cannot be known; therefore, we considered a theoretical maximum observable level of bycatch in each year as the basis for our simulations. Given the motivation for the enhanced datasheets and their design to provide more complete information about fulmar bycatch, the observation that enhanced datasheets could document greater numbers of fulmar bycatch in a year than the ASO database (2018; see Results), as well as the knowledge that enhanced datasheet reporting effort was highly variable and often less than ASO database reporting effort, we assumed that enhanced datasheets were more effective at documenting fulmar bycatch than the ASO database when given the same reporting effort. For the sake of (conservatively) assuming lower levels of both overall bycatch and bycatch mis-/under-reporting in the ASO database, we assumed that the high number of bycaught fulmars documented using enhanced datasheets in 2018 (114 birds) was a rare occurrence of the enhanced datasheets reporting the absolute maximum observable bycatch. We assumed this was rare both in the sense that the maximum observable bycatch in our simulations would usually be lower than this (representing some fraction of the unknowable total bycatch), and that the enhanced datasheet documented values would rarely reach proportions approaching 100% of this maximum. Both the ASO database- and enhanced datasheet-reported bycatch levels were then sampled probabilistically from the simulated maximum observable bycatch in every year, each based on an expected proportion that varied over time, i.e. a proportion which itself captured temporal variation in ASO effort that is unequal between the two methods, as well as the expected difference in method efficacies.

Halibut fishery fishing effort, fulmar populations, and ASO observer effort should be variable over time; therefore, both the total fulmar bycatch and the maximum observable level of fulmar bycatch should also be variable and positively skewed over time. Maximum observable bycatch was sampled independently in each year from a negative binomial distribution. Given the conservative assumptions of the model described above, we expected that most years (80% in the simulations) would have a lower maximum observable bycatch than the 114 birds observed in 2018; we also assumed that 20% of years would have maximum observable bycatch levels lower than the lowest possible value observed between 2016, 2018, and 2019 (i.e. 42 birds observed in the ASO database in 2016, which was the lowest possible observable bycatch across those three years when conservatively assuming that one of the methods actually documented the total observable bycatch). Having used the negative binomial distribution probability mass function to find parameters that satisfied these constraints, we simulated maximum observable fulmar bycatch using a negative binomial with mean = 80.2 and dispersion parameter *k* = 3.33 (lower *k* values describe higher positive skew; Hilbe, 2011).

Separate from year-to-year variation in fishery fishing effort and coverage, we also expected that ASO observation effort and coverage were not constant over time. The expected proportion of the maximum observable bycatch reported in the ASO database in each year was sampled from a beta distribution such that in 95% of years the documented proportion would be greater than what was observed in 2018 (10 fulmar documented in the ASO database compared to 114 from enhanced datasheets; i.e. a proportion of 0.0877, again assuming 114 was the maximum observable bycatch in that year). Additionally, despite having observed the very low “efficacy” of the ASO database in 2018, we still assumed that at least 1/3 of years would achieve greater proportions of the maximum observable bycatch than 50%. Using the beta distribution probability density function, we chose a beta distribution with α = 1.780 and β = 2.595 to satisfy these constraints (Ferrari and Cribari-Neto, 2004). Reported ASO database values were then sampled as binomial draws from the maximum observable bycatch using the expected proportion for each year.

We know that observer coverage for the enhanced datasheets should often be lower than that for the ASO database, and variable from year to year. To be conservative, despite the expected higher efficacy of the enhanced datasheets, we still parameterized the model such that expected proportions of the maximum observable bycatch detected using the enhanced datasheets would be less than that for the ASO database in 4/5 years. However, because we observed (presumably, for the sake of conservative model assumptions) one year out of three where the enhanced datasheets detected the maximum observable bycatch (2018), the simulated expected proportion has a 1/15 chance of being sampled from a uniform distribution between 90% and 100%. In other words, we assume that what we observed in one of three years was in fact a much rarer occurrence. In all other years (14/15), the proportion is sampled from a beta distribution with α = 1.200 and β = 6.400; this satisfies the overall condition that proportions should be less than those for the ASO database in 4/5 years. As before, the reported enhanced datasheet values were then sampled as binomial draws from the maximum observable bycatch based on the expected proportions.

All simulations were programmed in R (version 4.2.2; R Core Team, 2022).

## Results

### Enhanced seabird bycatch data sheets

While we actively distributed the enhanced seabird datasheets in 2016, 2017, and 2018, we received completed, enhanced seabird bycatch collection datasheets from the ASO companies in 2016, 2018 and 2019 from 0AB (Table 1). We did not distribute the enhanced seabird bycatch datasheets in 2019, but the ASO companies must have had some in the observer packages, and the ASO continued to complete them as requested by this project.

**Table 1.** Summary of northern fulmars (*Fulmarus glacialis*) and other birds reported in the enhanced seabird bycatch datasheets in three years of data collections with partners in the Greenland halibut (*Reinhardtius hippoglossoides*) fishery in North Atlantic Fishery Organization (NAFO) 0AB in northern Canada.

|  |  | Fulmars | Other birds | Total |
| --- | --- | --- | --- | --- |
| 2016 |  |  |  |  |
|  | Interacting with the gear, but not caught | 2 | 0 | 2 |
|  | Caught in gear, released alive | 24 | 0 | 24 |
|  | Major injury, but not caught | 0 | 0 | 0 |
|  | Dead in the gear | 25 | 10 | 35 |
| 2018 |  |  |  |  |
|  | Interacting with the gear, but not caught | 0 | 0 | 0 |
|  | Caught in gear, released alive | 0 | 0 | 0 |
|  | Major injury, but not caught | 0 | 0 | 0 |
|  | Dead in the gear | 114 | 0 | 114 |
| 2019 |  |  |  |  |
|  | Interacting with the gear, but not caught | 0 | 0 | 0 |
|  | Caught in gear, released alive | 6 | 0 | 6 |
|  | Major injury, but not caught | 0 | 0 | 0 |
|  | Dead in the gear | 17 | 3 | 20 |
| 2023 |  |  |  |  |
|  | Interacting with the gear, but not caught | 0 | 0 | 0 |
|  | Caught in gear, released alive | 0 | 0 | 0 |
|  | Major injury, but not caught | 0 | 0 | 0 |
|  | Dead in the gear | 2* | 6 | 8 |

\* Reported as 'gulls and terns' but confirmed as northern fulmars through photographs submitted with the datasheets.

In 2016 one ASO company submitted enhanced seabird bycatch datasheets directly to one of the authors of this paper. The reports came from two boats fishing for Greenland halibut in the 0AB region (all records were given as 0AB, and did not specify further). Observations of seabird bycatch were recorded between 10 July (first vessel with kits) and 29 October 2016 (last vessel with kits). Twenty-five fulmars were reported dead in the gear, with an additional 10 birds of unknown species reported dead (Table 1). An additional two fulmars were reported interacting with the gear, but not caught, and another 24 fulmars were reported caught in the gear but released in apparently healthy condition (Table 1). There were no reports of birds that interacted with gear resulting in major injuries and released without being brought onboard. All reports of bird bycatch were from baited gillnets.

In 2018, enhanced data collection sheets were only submitted through the carcass collection program and were submitted directly to the authors as part of that program (the carcass collection is described more below). Seabird bycatch was recorded between 13 June and 3 November 2018, and was reported as 0A, 0B or NA. From the carcass collection datasheets that included some enhanced seabird bycatch datasheets, a total of 114 fulmars were reported dead as seabird bycatch, but not necessarily brought onboard (Table 1). One datasheet in particular reported 100 northern fulmars as bycatch in a single set of a baited gillnets, with the note indicating “nets baited more than usual to use up remaining bait aboard”. Five bird carcasses from this set of 100 were submitted, all of which were confirmed as northern fulmars. Across all the submitted enhanced datasheets (*n*=114), a total of 113 of the fulmars were reported to have died in gillnets, with one individual fulmar having no gear type reported. Of these 113 fulmars, 110 fulmars were reported to have died in baited gillnets. There were no reports of birds that interacted with gear resulting in major injuries and released without being brought onboard.

In 2019, enhanced seabird bycatch datasheets were submitted by two ASO companies directly to DFO, which were then sent to the authors. Observations of seabird bycatch were conducted between 4 August to 18 September 2019, and all records were given as 0AB, and did not specify further. Three enhanced seabird bycatch datasheets were submitted from two different vessels in the fishery. Twenty birds were reported dead (but not necessarily brought aboard), of which 17 were recorded as fulmars and three were unknown bird species. There were an additional six fulmars caught in the gear and released alive. There were no reports of birds that interacted with gear resulting in major injuries and released without being brought onboard. All reports of seabird bycatch in 2019 were from baited gillnets.

In 2023, enhanced seabird bycatch datasheets were submitted by one ASO company directly to DFO, which were then sent to the authors. These datasheets referred directly to photographs that were taken of the birds reported that were submitted alongside the enhanced seabird bycatch datasheets. Observations of seabird bycatch were conducted between 10 to 29 October 2023, and all records were given as 0AB, and did not specify further. Two enhanced seabird bycatch datasheets were submitted from one vessel in the fishery. Eight birds were reported dead (some of which were brought onboard, some of which were lost before retrieval; Table 1). All of these were reported as ‘terns and gulls’ without more specific species level identification. For these enhanced seabird datasheets, two of these occurrences had associated photographs of the birds submitted. Examination of those photographs showed that at least two of the birds reported as ‘terns and gulls’ were northern fulmars (Figure 2). There were no reports of birds that interacted with gear resulting in major injuries and released without being brought onboard. All reports of seabird bycatch in 2023 were from baited gillnets.

**Figure 2.**
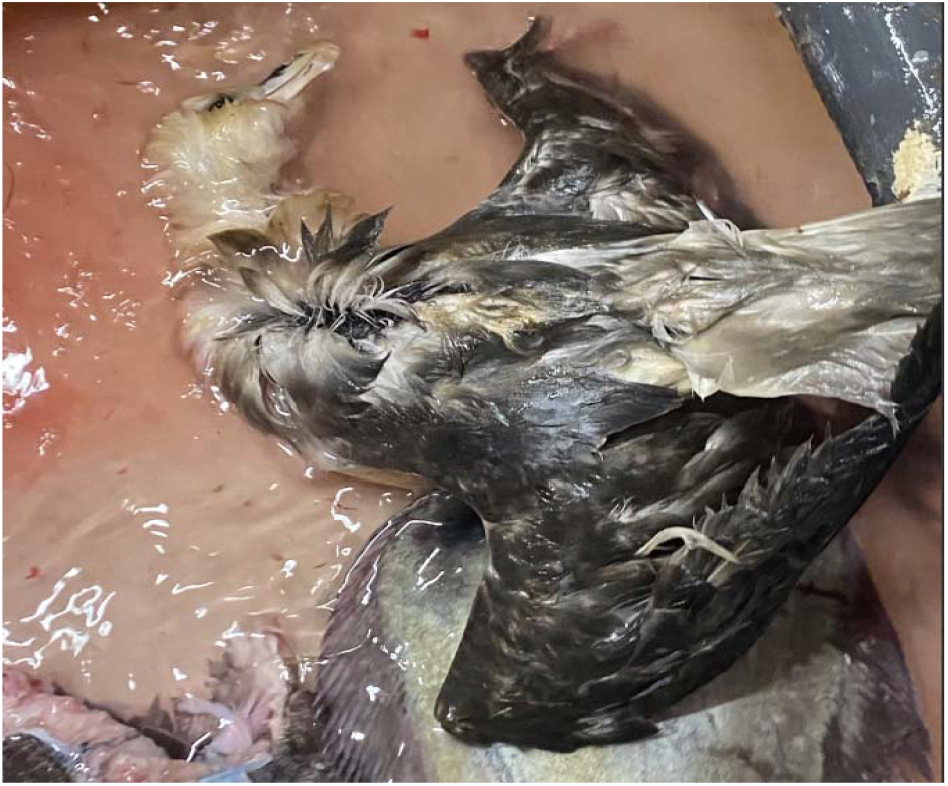
A northern fulmar (*Fulmarus glacialis*) reported as incidental catch in the Greenland halibut (*Reinhardtius hippoglossoides*) fishery in NAFO 0AB in northern Canada, but which was reported in the ‘tern and gull’ category by the At-sea Observer (ASO) and submitted as part of the data collection for this project.

### Carcass collections

As previously reported (Colston-Nepali et al. 2020), 19 seabirds from the 2018 fishing efforts in 0AB were submitted to ECCC, but only seven of these were identified as fulmars on the datasheets (the other 12 listed as unknown species) (Figure 1C). Visual inspection and genetic techniques subsequently confirmed that all 19 birds submitted were fulmars (Colston-Nepali et al. 2020).

### Comparison across data collection methods

For three years of our study we received data from two or more methods of reporting seabird bycatch. The number of fulmars and total birds reported as bycatch in the 0AB Greenland halibut fishery in 2016, 2018, and 2019 differed among the data collection methods (Figure 3). In 2016 a total of 42 fulmars were reported in the DFO ASO database, while fewer (25 fulmars and 10 unknown birds) were collected via the enhanced seabird data sheets. In 2018, 10 fulmars were reported in the DFO ASO database, while more (19 fulmars) were submitted via the carcass collection programs, and far more (114 fulmars) were recorded on the enhanced seabird collection datasheets. In 2019, 44 seabirds were reported in the DFO ASO database, while fewer (17 fulmars and 3 unknown birds) were collected via the enhanced seabird data sheets.

**Figure 3.**
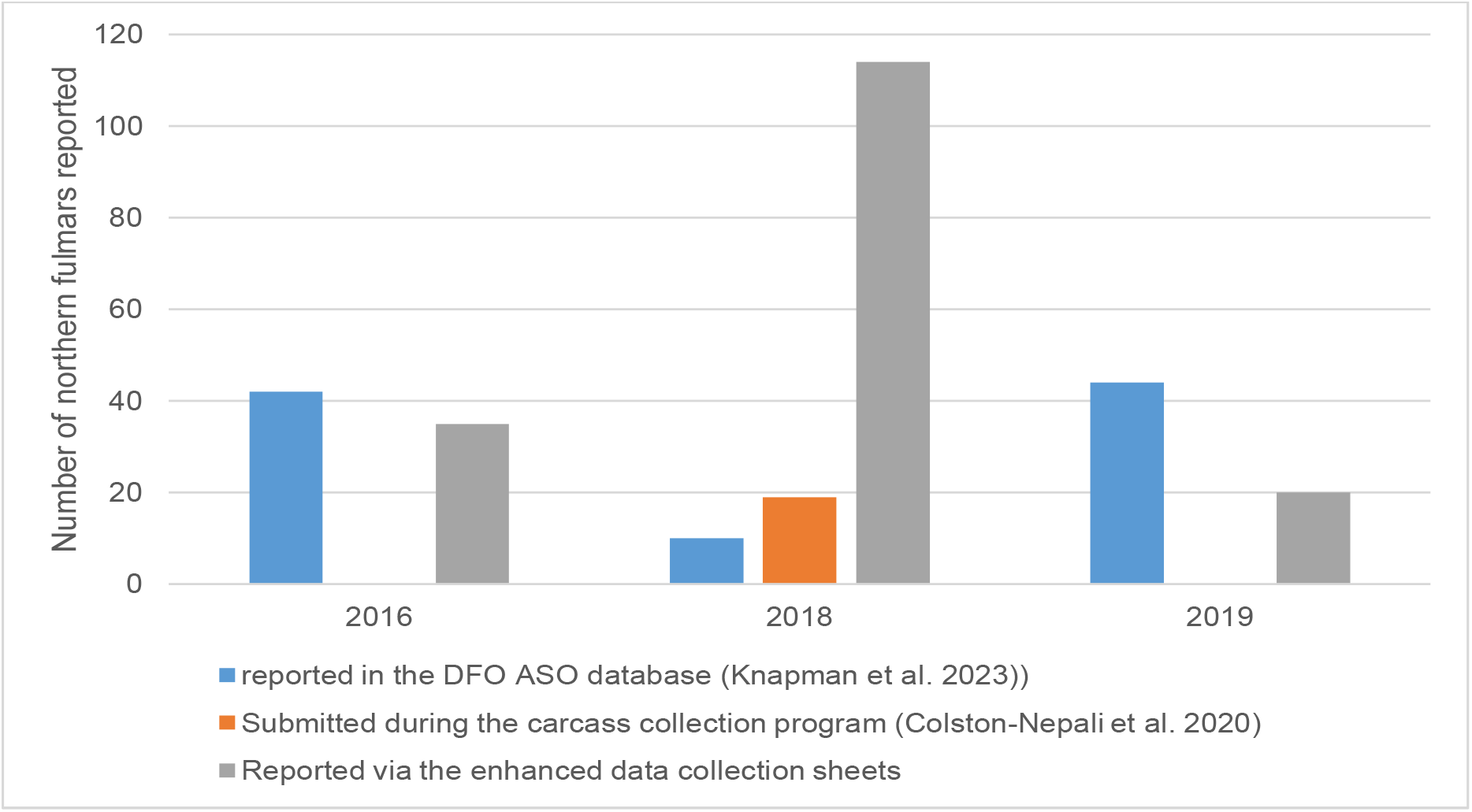
Numbers of northern fulmars (*Fulmarus glacialis*) and other birds reported different seabird bycatch collection methods across three years of data collections with partners in the Greenland halibut (*Reinhardtius hippoglossoides*) fishery in NAFO 0AB in northern Canada.

### Simulation

The output from the simulation model is presented in Figure 4 and demonstrates how the assumed drivers of observed variation in detected fulmar bycatch produces reported values that are inconsistent in their relationship with the hypothetical maximum observable level of bycatch, regardless of observation method. While the simulated ASO database records are often higher than the enhanced database values, the former are often still much lower than the maximum observable bycatch. If variation in the maximum observable bycatch is largely linked to variation in actual bycatch, ASO records will at best only weakly reflect temporal trends in total bycatch levels. Enhanced datasheets often reported only small proportions of the observable maximum in the simulations, though occasionally – as was observed in 2018 – greatly outperformed the ASO database detections. These simulated values demonstrate how, based on our interpretations of the observed data, neither ASO database records nor enhanced datasheet records can serve as reliable indicators of Greenland halibut fishery 0AB fulmar bycatch in any given year, or as indicators of temporal trends in that bycatch, given current inconstant observation methods, efforts, and coverage. Note that had we not made some generous assumptions about the ASO database records (e.g. that the relative efficacy observed in one of three years was a 1/15 occurrence; that the enhanced datasheets should be expected to detect lower proportions of bycatch than the ASO database in 4/5 years) the simulations would only have indicated worse discrepancies between the ASO database records and the maximum observable bycatch.

**Figure 4.**
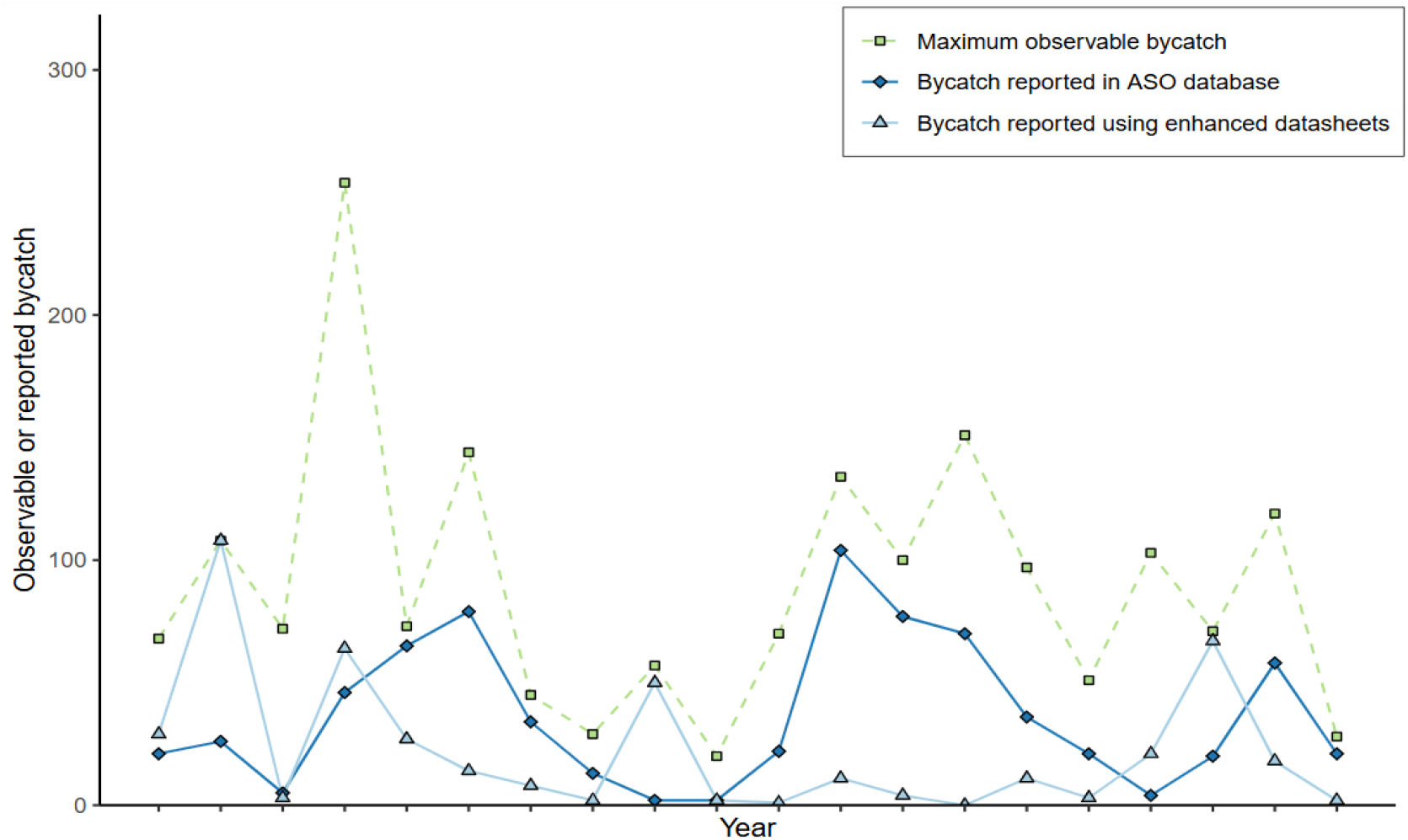
Simulated maximum observable bycatch, bycatch reported in the ASO database, and bycatch reported using the enhanced datasheets across 20 years.

## Discussion

Acquiring accurate detailed data for seabird bycatch is an ongoing challenge to effective implementation of an ecosystem approach to managing fisheries (Anderson et al. 2011; Regular et al. 2013; Jiménez et al. 2014; Christensen-Dalsgaard et al. 2019). This is especially true in the Arctic region where different countries have highly variable reporting and data systems to track and understand seabird bycatch in fisheries (Bærum et al. 2019; Christensen-Dalsgaard et al. 2019; O’Keefe et al. 2023). In this study, we found that employing different methods for the data collection of seabird bycatch in the NAFO 0AB fisheries produced markedly different results in species reported and total numbers of birds reported, which has implications on how any future work may model seabird bycatch as a threat to sustainable seabird populations. Widely disparate numbers between reported and real bycatch have also been found in the Baltic Sea (Morkunas et al. 2022), and this study adds to our further understanding of the nuances of global fisheries bycatch data.

While other non-fulmar species are reported in the datasheets collected as part of this project (e.g., 2019), to date we believe that all birds bycaught in 0AB are northern fulmars. This is based on previous studies examining the other species in the ASO database and their global ranges and other data sources (Anderson et al. 2018), and on the visual confirmation from photographs (reported here from our 2023 data), and using genetic confirmation of fulmars (Colston-Nepali et al. 2020). Therefore, we feel confident assuming the total number of birds reported here are fulmars. Thus, while only 17 fulmars were reported in 2019, it is highly likely that all 20 birds reported were fulmars. Given that 44 fulmars were reported in the ASO database that year, this finding fits with our predictions that the total counts of fulmars would be lower (by 45% in 2019) in these enhanced methods, because these methods would not be applied universally across the vessels and ASO. Similarly, in 2016 we found 35 birds using the enhanced data collection sheets, but 42 were reported in the ASO database, suggesting that 83% of the birds were reported in both methods. This demonstrates that while the ASO were asked to complete the enhanced seabird bycatch collection sheets, not all the ASO completed this task as expected, and provided a subset of more detailed data as compared to the ASO database. These findings suggest that the expected nestedness of these data types is less perfect (Figure 5A).

**Figure 5.**
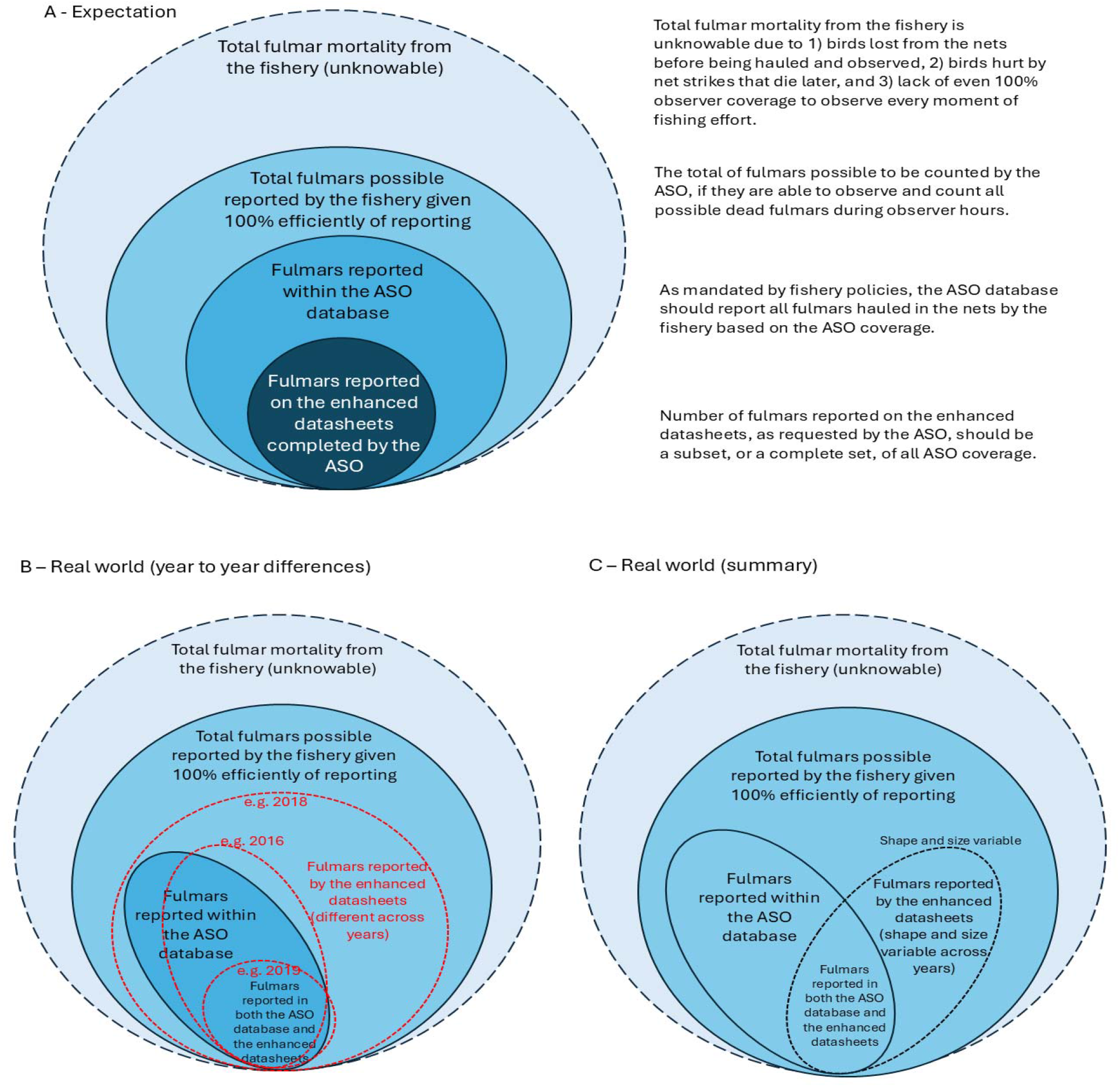
Schematic representations of nestedness of seabird bycatch reporting, and real reports. A – expected levels of bycatch across data types, B – observed differences in the fulmar bycatch in the different data types in different years, and C – summary of how the different data types vary in relation to each other year to year, generally.

The most notable difference in total individual fulmars reported was in 2018 where the ASO reported a total of 10 fulmars (and no additional birds reported), while the carcass collection program yielded 19 fulmars, almost twice as many individuals. Even more surprising, the 2018 enhanced seabird bycatch datasheets report 114 fulmars – approximately 11X the number of bycaught fulmars reported in the ASO database that year. This unexpected finding suggests that in any given year the total fulmar numbers reported in the ASO database may be inaccurate by an order of magnitude, even in areas that should have close to total coverage by ASO (Figure 5B; the enhanced seabird datasheet scenario for 2018). This finding has implications for any project looking to model what the potential impacts of this biological removal of fulmars by the fishery are on the fulmar population (needed to ensure ancillary effects of the fishery do not affect the sustainability of sympatric wildlife populations), because these inaccuracies are critical to consider. For example, Anderson et al. (2018) modeled “estimated current” fulmar bycatch in this fishery and a “10X” model with 10X the current estimated bycatch; the “current” model suggested a small, long-term decline in numbers, whereas the “10X” model led to rapid, catastrophic declines in fulmar numbers.

In 2019, we also found that 90% of the birds that had interactions with fisheries gear were reported as dead (although we cannot ascertain if these are the same individual birds as in the ASO database). This is also an important finding because population modelling exercises (e.g., Anderson et al. 2018) using ASO data need to consider those birds that have interactions with gear, and may be likely to die post-release, but are not taken onboard so not reported in the ASO database (Brothers et al. 2010). This is important to consider as it is a contributing factor to knowing the true level of mortality of any fishery (Figure 5A). In 2016, while 35 birds were reported dead, and additional 24 fulmars were caught in the gear and released alive. If most or all of those released birds died, then using ASO data to model bycatch mortality would underestimate bycatch mortality by ∼40%.

The sources of these varying values in seabird bycatch numbers among the methods are not surprising given the different methods we report, but the magnitude of the differences is concerning, and the source of the variation is important to consider. All of the data considered here – the traditional ASO database, the enhanced seabird bycatch datasheets, and the carcass collections – came from the same ASO effort onboard the vessels (we have omitted ASO IDs and other identifiable information to anonymize the data) (Figure 5). Thus, these data should be nested subsets of each other (Figure 5A), but the data presented above illustrate that they are related but the degree of overlap of the data differs among years (Figure 5B) resulting in an unknown relationship between the datasets that is highly variable (Figure 5C). This led to the relatively large differences between the maximum observable bycatch levels, and the reported values in the ASO database and the enhanced datasheets. In turn, this suggests that this type of variability within the known bycatch data sources needs to be considered within any population effect assessments in order to more accurately represent the number of individual fulmars that are removed from the population in any given year.

Importantly, while not all the birds reported were attributed to a NAFO division they were bycaught in beyond 0AB, 107 of the 114 fulmars reported in 2018 were from NAFO Division 0A, which for the target fishery in this study has targeted 100% ASO coverage for gillnets (the only gear type we had reports from in this study). The traditional ASO database is the requirement for the ASO as part of their duties, and the voluntary enhanced seabird bycatch datasheets as well as the carcass collections were in addition to those duties. Therefore, the data presented in each method was a nested set of data that counts the same seabirds bycaught and represents how comparable those data methods are to each other within a year. While the ASO coverage varied among NAFO divisions, and likely across years, all the data from this work were from the ASO effort, which was a constant in any given year. Hence, our findings suggest that the traditional ASO observer database protocols and data collection are underestimating the total fulmars bycaught in the fishery in at least some years.

This work is particularly important when considering the ecosystem impact of fisheries within the context of marine stewardship. Currently, the Greenland halibut fishery has been certified by the Marine Stewardship Council (MSC) (Knapman et al. 2019). The certification final report included several conditions around seabird bycatch due to previous work showing the potential impacts on fulmars from the fishery (Anderson et al. 2018; Knapman et al. 2019). Specifically, the certification stated that:

> “By the fourth surveillance audit the client shall provide evidence that there is a partial strategy in place, if necessary, for the UoA (Unit of Assessment) that is expected to maintain or not hinder rebuilding of northern fulmar at/to levels which are highly likely to be within biologically based limits or to ensure that the UoA does not hinder their recovery” (Knapman et al. 2019:72).

Therefore, updated and rigorous data are required on which to determine whether these conditions have been met during the surveillance and renewal cycle, and all other fisheries management practices (DFO 2007).

Accurate data collection on seabird bycatch is also critically important to inform co-management in the region, a cornerstone of Arctic wildlife management (Armitage et al. 2011). In northern Canada where this fishery takes place, fisheries are managed by DFO and the Nunavut Management Wildlife Board, as directed under the land claim agreement for Nunavut (Daoust et al., 2010). However, both organizations also have responsibilities for managing seabirds as they relate to fisheries activity (along with ECCC). Therefore, detailed information on both the effort for the target species, as well as detailed information about all bycatch species are needed for those organizations to make evidence-based policy decisions (Votier et al. 2023).

Within the context of these data, it is important to consider that although DFO policies indicate that seabird bycatch data should be collected by fisheries (DFO 2019a; DFO 2019b), the collection of seabird bycatch data is not currently an explicit condition of the license in the Greenland halibut fishery in 0AB (C. Friesen, DFO (Winnipeg), pers. commun., April 2024). Therefore, when examining any data produced under various methodologies in this fishery, including interannual variation of bycatch, total bycatch values, or the effectiveness of mitigation, the data only present a partial picture in any given year as the collection of the data is unlikely to receive similar effort across years. To improve consistent data collection on seabird bycatch, a condition of DFO license that includes seabirds and specifically lists seabird species should be implemented.

We note that the data from 2016, 2018, and 2019 were collected before the MSC process that the fishery undertook for sustainability certification (Knapman et al. 2019). Following the implementation of the MSC certification process and the resulting condition specifically mentioning northern fulmar bycatch as a potentially problematic bycatch species, actions have been taken by the industry to minimize seabird bycatch. Therefore, future work should be directed towards analyzing data on an ongoing and annual basis to consider the effects of these changes on bycatch levels.

A limitation with this work is that it is difficult to extrapolate these findings as there is no way to measure effort in relation to these reported numbers (as outlined in Figure 5). This also limits any statistical modeling we can carry out as we do not have a true value against which we can evaluate these methods. We cannot assess whether these results apply across the fleet, or to a certain subset of vessels. The reports for some birds also identified the fishing region as 0AB, and did not differentiate between 0A and 0B, which had differing ASO coverage. Some studies have used both seabird bycatch and fishing effort to extrapolate how many birds are taken at the fishery level (e.g., Waugh et al. 2008; Phillips 2013; Christensen-Dalsgaard et al. 2019). In the Division 0AB fishery for Greenland halibut, because of the few vessels active in the fishery, the fishing effort data are not available. For example, as part of our overall work on seabird bycatch in the Canadian Arctic, we requested the fishery effort data for the Greenland halibut NAFO Division 0BA, but because of the Privacy Act in Canada and the “Rule of 5” around the release of data (Tomasic 2023), DFO was unable to share this information. Without being able to confirm which NAFO Division the birds are from within 0AB, or the associated effort data on the use of these enhanced methods to report seabird bycatch, all we can conclude is that the numbers presented here represent an absolute minimum for the fishery. Matching seabird bycatch data to the related effort data in Arctic Canada remains a challenge, even with these enhanced data collection efforts.

Importantly, while we requested the ASO companies to collect enhanced data on seabird bycatch in a variety of NAFO Divisions (NAFO Subareas 0-5) that their companies serviced, we only received data from ASO from the NAFO 0AB Greenland halibut fishery. Clearly, additional uptake and adoption of seabird bycatch monitoring and transparent reporting is required. This seems particularly pertinent for the Greenland halibut fishery in Arctic Canada where efforts to reduce fulmar bycatch and demonstrate measures to promote sustainable requirements are a condition of MSC certification; the precautionary principle as applied to fisheries (González-Laxe 2005) suggests that current efforts are insufficient for robust and reliable modelling.

This work highlights how data on incidental seabird bycatch can vary among methods employed, and future work should consider expanding similar enhanced datasheet collection efforts to other regions to increase the robustness of data on seabird bycatch in Canada. To improve our ability to provide evidence for sustainable ecosystem level management, rigorous and consistent data collection, data management, and data analyses needs to be prioritized on an annual basis for incidental catch of seabirds. Given the within-year variability we recorded, evidence of declining Arctic fulmar numbers, and the absence of a robust seabird bycatch data collection plan, we strongly recommend caution in permitting increased fisheries effort in NAFO Divisions 0AB for gear types that seabirds are known to be susceptible (e.g., gillnets and longlines) until the effects of this fishery on local seabird numbers can be better addressed.

## Author Contributions

JP conceived of the ideas and designed the methodology. JP collected, curated and analysed the data, while JP and MM made interpretations of information. AM designed and implemented the modeling aspects of the paper. JP, AM and MM wrote the paper, contributed critically to manuscript drafts, and gave approval for publication.

### Acknowledgements

We appreciate the ongoing collaboration with the Nunavut Fisheries Association, as without it, this work would not have been possible. Partners at the DFO were also instrumental in setting up this program to ensure collection methods employed aligned with other methods used in the fishery. We thank the fishers, crew members, and at-sea observers that put in the extra effort to collect these data as part of this project. We thank colleagues at DFO, Oceans North, and ECCC for feedback on initial drafts of the work, and *** reviewers for their comments which helped to improve the manuscript.

## Conflict of Interest Statement

The authors declare no conflict of interest with this work.

## Supplemental Material

**Figure A.**
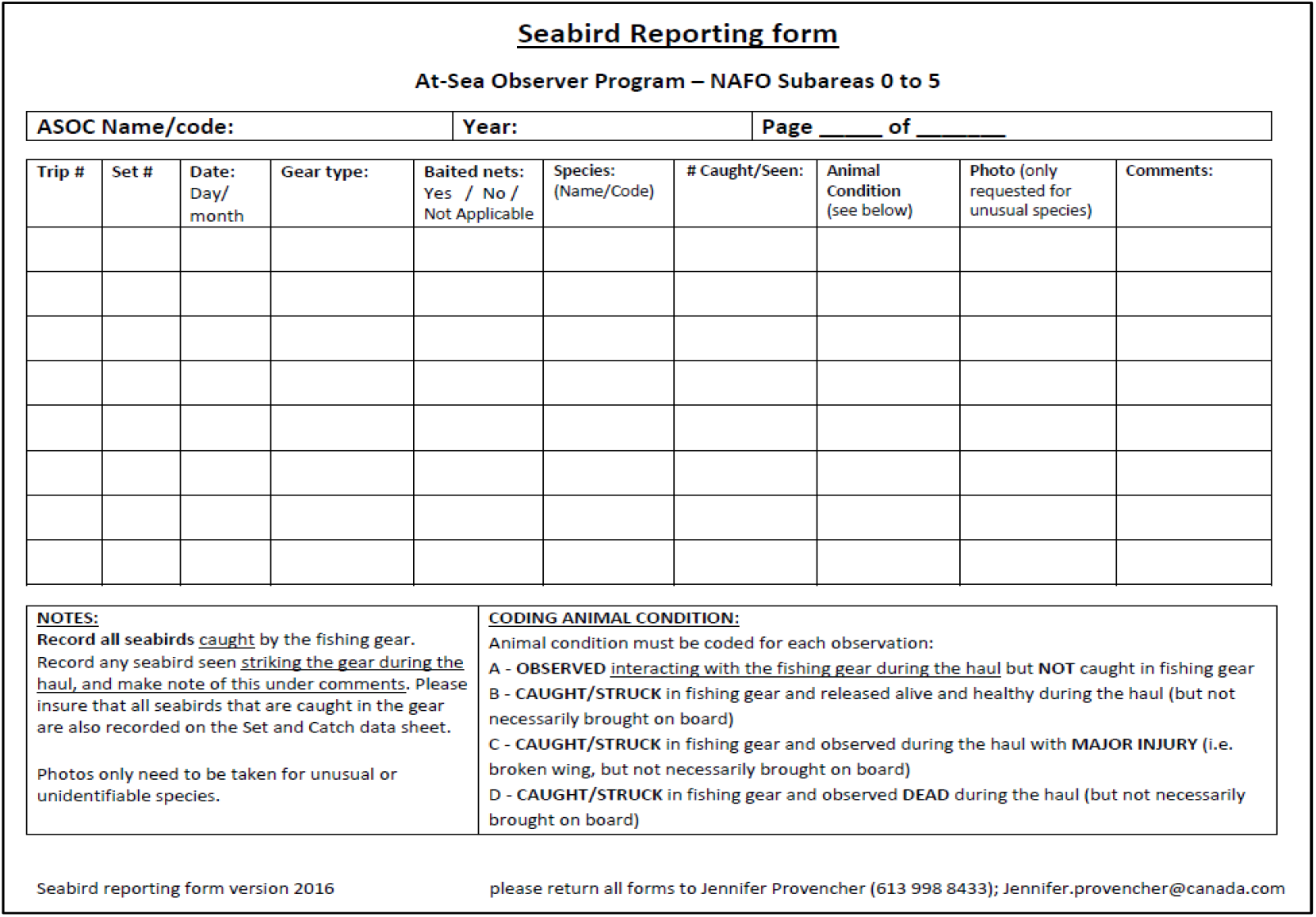
Enhanced incidental seabird bycatch datasheet distributed to the At-sea observers (ASO) issued in 2016 as part of the project to collect more detailed information about seabird interactions with fisheries gear in the NAFO 0AB region.

**Table A.** Summary of northern fulmars (*Fulmarus glacialis*) and other birds reported different incidental seabird bycatch collection methods across three years of data collections with partners in the Greenland Halibut (*Reinhardtius hippoglossoides*) fishery in NAFO 0AB in northern Canada.

|  | 2016 | 2018 | 2019 |
| --- | --- | --- | --- |
| reported in the DFO ASO database (from Anholt et al. in (Knapman et al. 2023)) | 42 fulmars | 10 fulmars | 44 fulmars |
| Submitted during the carcass collection program (Colston-Nepali et al. 2020) | NA | 19 fulmars | NA |
| Reported via the enhanced data collection sheets | 25 fulmars,<br>10 unknown bird species | 114 fulmars | 17 fulmars,<br>3 unknown bird species |

## Notes

### Competing Interest Statement

The authors have declared no competing interest.

